# Homogenization of a Reaction Diffusion Equation can Explain Influenza A Virus Load Data

**DOI:** 10.1101/2021.01.04.425332

**Authors:** Arwa Abdulla Baabdulla, Hesung Now, Ju An Park, Woo-Jong Kim, Sungjune Jung, Joo-Yeon Yoo, Thomas Hillen

## Abstract

We study the influence of spatial heterogeneity on the antiviral activity of mouse embryonic fibroblasts (MEF) infected with influenza A. MEF of type *Ube*1*L*^−/−^ are composed of two distinct sub-populations, the strong type that sustains a strong viral infection and the weak type, sustaining a weak viral load. We present new data on the virus load infection of *Ube*1*L*^−/−^, which have been micro-printed in a checker board pattern of different sizes of the inner squares. Surprisingly, the total viral load at one day after inoculation significantly depends on the sizes of the inner squares. We explain this observation by using a reaction diffusion model and we show that mathematical homogenization can explain the observed inhomogeneities. If the individual patches are large, then the growth rate and the carrying capacity will be the arithmetic means of the patches. For finer and finer patches the average growth rate is still the arithmetic mean, however, the carrying capacity uses the harmonic mean. While fitting the PDE to the experimental data, we also predict that a discrepancy in virus load would be unobservable after only half a day. Furthermore, we predict the viral load in different inner squares that had not been measured in our experiment and the travelling distance the virions can reach after one day.

## 1. Introduction

In experiments of mouse embryonic fibroblasts (MEF), we infected MEF of type *Ube*1*L*^−/−^ with influenza A virus to observe their susceptibility and resistance to viral infection. The fibroblast population is bimodal, consisting of cells that sustain a strong or weak virus infection, respectively (Now and Yoo, 2016). In typical cell experiments, these cells are mixed, forming a homogeneous population. The importance and correlation between the virus infection and population with heterotypic patterns has been studied in (Snijder et al., 2009). However, the effects of spatial complexity between heterogeneous populations has not been explored yet. Therefore, it is of interest to understand whether the viral susceptibility changes when the spatial distribution of the sub-populations becomes heterogeneous.

To analyse this question, (Park et al., 2017) employed a brand new cell-printing method, which allowed printing of cell cultures in a checker board pattern, where the two cell populations are separated into little squares (see Figure 1 (A)). The size of the inner squares can be adjusted from almost complete separation (large squares), to finely mixed (small squares), to fully mixed. In Park et al. (Park et al., 2017), the experimental plates had 50% of A549 human alveolar lung epithelial cells, and 50% of HeLa cervical cancer cells. These two cell lines are known to possess relatively “strong” and “weak” infectivity to influenza A, respectively (Li et al., 2009; De Vries et al., 2011). To their surprise, the total virus load did *“ not arise as a simple arithmetic summation of the individual cellular activities”*. Here we repeat the experiments with mouse embryonic fibroblasts (MEF) of type *Ube*1*L*^−/−^ using 50% “weak infectivity” sub-type and and 50% “strong infectivity” sub-type. Similar to the experiments of (Park et al., 2017) we also find that the total virus load after one day of inoculation depends on the spatial arrangement of the cells. The finely mixed and fully mixed plates had a much reduced total viral load as compared to the large scale patterns (see data in Figure 1 (B)), suggesting a strong dependence of the total viral load on the spatial arrangement.

**Figure 1:**
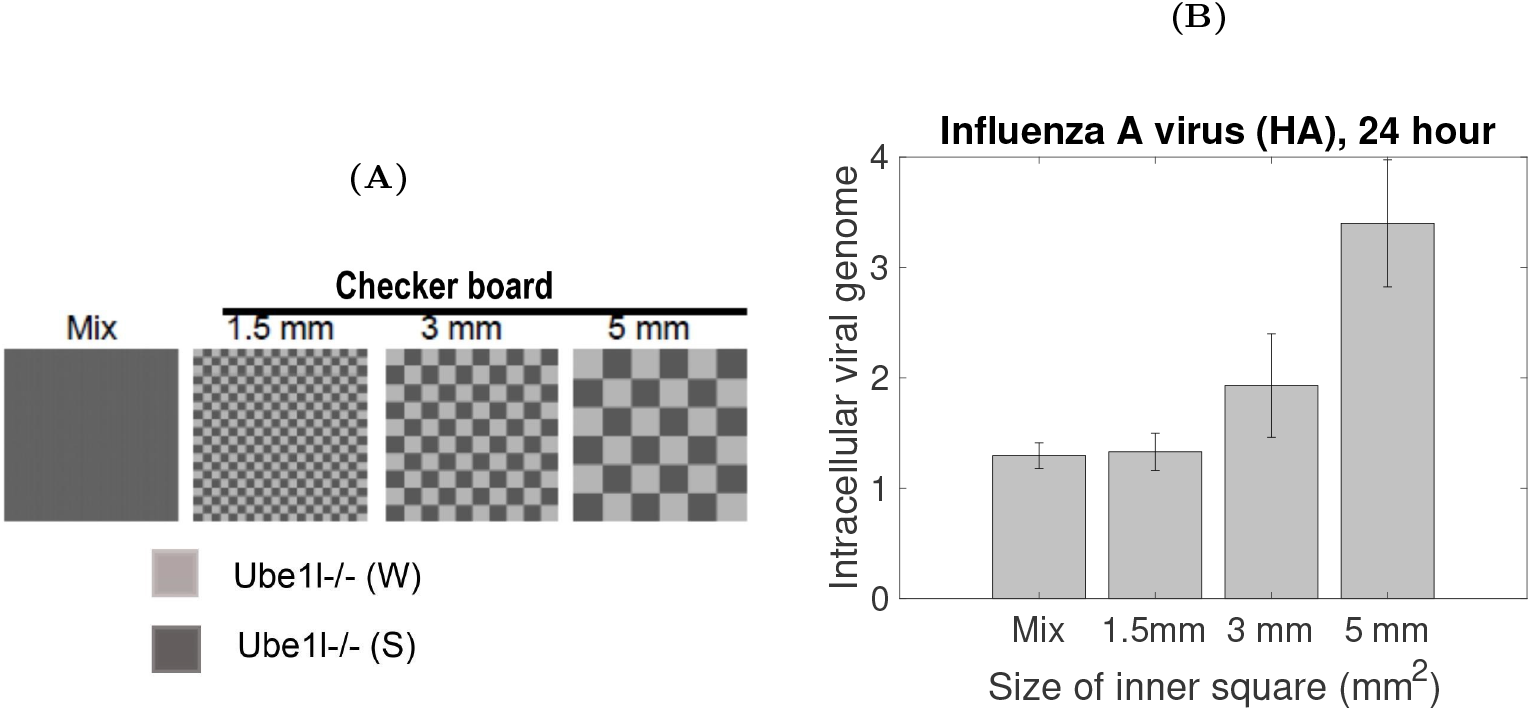
**(A):** Design of cell-printing pattern, **(B):** Viral load measurement after 24h.

In this paper we use mathematical modelling to explain the spatial dependence of the virus load data. Mathematical modelling of virus load is an active field of research (see Smith and Perelson (2011); Beauchemin and Handel (2011); Gallagher et al. (2018); Smith (2018)). While most of these models are based on ordinary differential equations, a few approaches use spatial modelling (Gallagher et al. (2018); Wodarz et al. (2012); de Rioja et al. (2016)). Here we base our model on a Fisher-KPP reaction-diffusion model (Murray, 2007) and we use homogenization methods and numerical methods to analyze the model. To say it upfront, while going from coarse to fine mixture, we observe a transition from an arithmetic mean of two steady state virus load levels to a harmonic mean of these values. As the harmonic mean is always smaller than the arithmetic mean, it explains the observed reduction in virus load for the finer mixed experiments. As we calibrate the model to our experimental data we find that the model can not only explain the observed virus load data, it can also predict values that were not measured in the experiments. Moreover, we found the spatial viral load distribution across the periodic domain by solving the mathematical model numerically.

Our analysis shows a simple but relevant example of the effect of spatial heterogeneity on cell responses to virus infection. It shows that measurements done on cell populations in isolation cannot simply be carried over to a heterogeneous mixture of cells. The spatial arrangement seems to be important.

### 1.1. Outline

The paper is organized as follows: In the next section (Section 2) we explain the experimental set up and the data collection. In Section 3 we introduce the mathematical model. We chose a very classical Fisher-KPP reaction-diffusion equation (Murray, 2007), which is quite sufficient for our purpose. We then perform the spatial homogenization as it is relevant for our problem. From this analysis the dichotomy between arithmetic and harmonic means arises. In Section 4 we fit our model to our virus-load data using a log-likelihood method, thereby explaining the observed virus load dependence on the spatial arrangement. In Section 5 we present numerical solutions of the corresponding model, which show the spatial distribution of the viral infection across the checker board pattern. We close with a discussion in section 6.

## 2. Influenza A Infection Experiments

We consider mouse embryonic fibroblast (MEF) of *Ube*1*L*^−/−^, where *Ube*1*L* stands for a ubiquitin like modifier activating enzyme for ISGylation protein that conjugates an Interferon (IFN) stimulated gene 15 (ISG15) to target proteins. The *Ube*1*L*^−/−^ are null mutations of *Ueb*1*L*, where the *Ueb*1*L* production is deactivated. Studying these cells, we found that *Ube*1*L*^−/−^ populations are bimodal, with two sub-populations of differential antiviral activity (Now and Yoo, 2016). *Ube*1*L*^−/−^(*S*) and *Ube*1*L*^−/−^(*W*) designates those subpopulations with *strong* and *weak* infectivity, respectively. The viral infection data for each population in isolation are listed in the Appendix A in Tables A.5 and A.6.

To study the influence of the spatial distribution patterns on the virus load of the population as a whole, we printed *Ube*1*L*^−/−^(*S*) and *Ube*1*L*^−/−^(*W*) cells in a regular checker board pattern by using the inkjet bio-printing system (Park et al., 2017). While fixing the size of the checker board square to 30 × 30 *mm*^2^, the size of the inner squares was varied using side lengths of 1.5 mm, 3 mm, 5 mm, and fully mixed, (see Figure 1 (A)). Thus, the geometric separation of *Ube*1*L*^−/−^(*S*) and *Ube*1*L*^−/−^(*W*) cells is increased as the size of the inner squares is increased. The mixed case of 50/50 of *Ube*1*L*^−/−^(*S*) and *Ube*1*L*^−/−^(*W*) cells is used as a control group. In each of these experiments, the cells on the checker board and the mixed plate were infected with Influenza A virus and incubated for 24 hours. The inoculation was performed unifromly over the entire domain to avoid spatial heterogeneities through the inoculation process. The infected cells were harvested and the total amount of intracellular viral genome was measured using the real time quantitative PCR method (Figure 1 (B)). All experiments use the inkjet printing method that was developed in Park et al. (2017), The experiments for the cell cultures in isolation were carried out twice for each cell type, and in each case three plates were inoculated with a density of about 6 10^6^ cells/mL. The homogeneous cell populations were printed with the same set up as for the 1.5 mm checker board printing, however, each square would carry the same cell type (all black or all white, respectively). The results for the homogeneous populations are reported in Tables A.5 and A.6. The checker board measurements were also repeated three times, and the results are shown in A.7.

The method of Livak (Delta Delta CT) has been used to compute the relative quantification (gene expression)

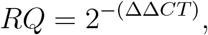

where *CT* represents the cycle number where the fluorescence that is generated by the PCR of the influenza A gene is distinguishable from the background noise cycle threshold (CT) of our sample. We measure Δ*CT* by the following formula

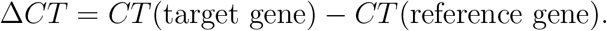

Here, our target gene is influenza A hemagglutinin (HA) gene and the reference gene is the mouse glyceraldehyde 3-phosphate dehydrogenase(mGAPDH) gene. Thus, we can compute ΔΔ*CT* as the following

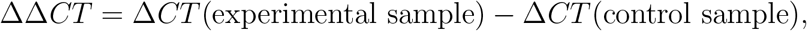

The full data set is shown in Appendix A.5 and A.6 for the homogeneous populations and in A.7 for the checker board data. The data from A.7 are shown in Figure 1 (B).

Looking at the virus load data in Figure 1 (B), we can see clearly a mismatch in the viral load depending on the inner square size from mix, to 1.5 mm, to 3 mm and to 5 mm. However, each experiment has the same ratio and the same mass of *Ube*1*L*^−/−^ (W) and (S) cells. Therefore, the antiviral activity of the cell population as whole is not a simple summation of the individual cellular activities.

## 3. A Mathematical Model

Reaction diffusion equations (RDE) are a powerful tool whenever the spatial spread of a population is of importance. One of the simplest examples of a RDE equation is the Fisher-KPP equation

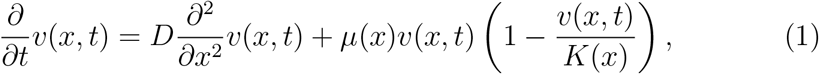

where *v*(*x, t*) is the viral density at time *t* and location *x*, *D* is the diffusion coefficient, describing the spatial spread of the virions, *μ* is the population growth rate of the virus, and *K* is the population carrying capacity. The viral infection and replication inside the cells is a multilayered processes. First, virions enter the cell through endocytosis. Then, after virion uncouating, the virus RNA replicates and is reassembled into new virions. Virions are released through a number of processes, such as exocytosis and cell lysis. In the case of lysis the cell membrane breaks down, releasing virions into the extracellular fluid. Biologically, the virion growth rate parameter *μ* in our model can be seen as an effective growth rate, combining the details of viral replication inside cells. While, the diffusion coefficient describes the free virions spread. Since the model is used to describe the spatially varying checker board patterns, we assume that the growth rate *μ*(*x*) and the virus carrying capacity *K*(*x*) are spatially dependent. We assume that the transport of virus from cell to cell is the same for all cell types, hence we assume *D* is not spatially dependent and it is constant. The model can be considered for the case of *D*(*x*) as well (see Shigesada et al. (1986, 2015); Maciel and Lutscher (2013)), but the model with constant *D* is sufficient to explain our data. Moreover, we have no biological indication to assume that the diffusivities should be different, hence we assume they are the same.

Fisher proposed equation (1) in his paper *“The wave of advance of advantageous genes”* in 1937 (Fisher, 1937). He studied the diffusion of species in one dimension and its traveling wave solutions with considering the reaction term being logistic. In the same year, Kolmogorov, Petrovsky, and Piskunov studied the reaction diffusion equation in two dimensions and with more general monostable reaction term (Kolmogorov et al., 1937).

We chose the Fisher-KPP equation for our modelling problem for several reasons. Firstly the available data on virus load on checker board patterns are limited to the total virus load at a few time points (0, 6h, 12h, 24h). No microscopic measurements are performed, hence no details on the virus replication inside cells, the cell bursting, number of released virions, transport of virions inside the cell tissue, and cell death are available. It would be fantastic to include those details in a more sophisticated modelling framework, but at this stage this is neither possible nor needed. We find that the simple Fisher-KPP approach is entirely sufficient to explain the phenomenon on a macroscopic level. A second reason to use Fisher-KPP is that it has proven useful in many applications before, it is simple, and the behavior of this model is well understood (Murray (2007)). There are certain model characteristics we can use immediately. For example the invasion speed of the Fisher-KPP model (1) is

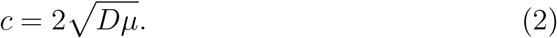

The parameters of the Fisher-KPP equation (1) *D, μ, K* will be estimated based on the data from the experiments which we presented in the previous Section 2.

### 3.1. Homogenized Fisher KPP Model

The experimental set up as described above is a paradigm for a homogenization problem. A microscopic scale (inner squares) is varied on a finer and finer scale, until in the limit, a homogeneous mixture arises. We are in the fortunate position, that not only the separated and fully mixed states are measured, but also several intermediate values for intermediate mixture types. While homogenization is a well known scaling method in physical applications (Pavliotis and Stuart, 2008; Holmes, 2012), it has only recenlty been used for ecological problems in (Garlick et al., 2011; Maciel and Lutscher, 2015; Yurk, 2018; Yurk and Cobbold, 2018). To our knowledge, this method has never been used in the microbiological context considered here.

Due to the symmetry of the problem, we present the argument in a one-dimensional setting. The scaling method applies to higher dimensions as well, but the one-dimensional setting is sufficient for our purpose. To model the specific checker board pattern, we divide the real line into small intervals of equal length, which separates weak and strong infectivity populations (see Figure 2). On this periodic domain we consider the spatial dependent Fischer-KPP equation (1), where the virus growth rate *μ*(*x*) and the virus load carrying capacity *K*(*x*) vary between cell types, i.e.

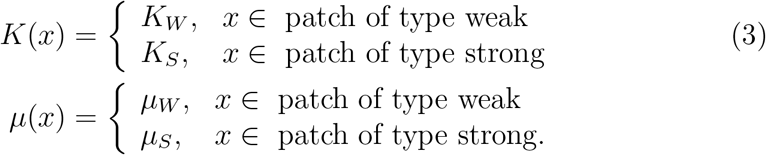

**Figure 2:**
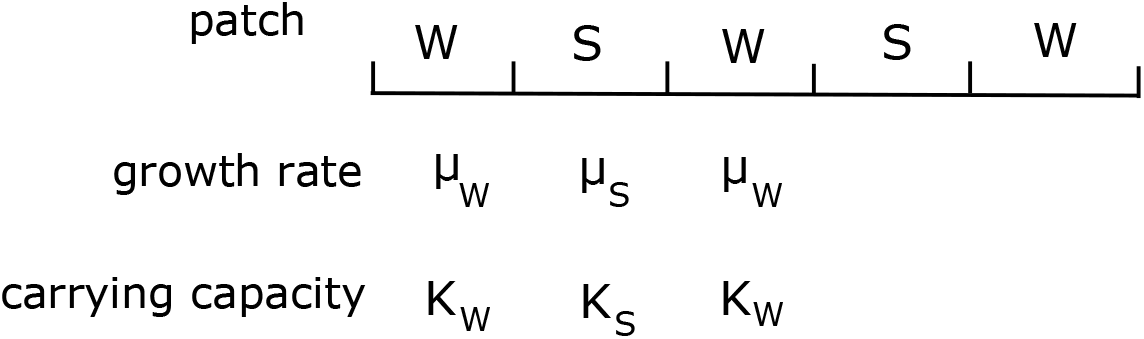
Sketch of a periodic patchy environment of two cell types.

We use W to indicate the *Ube*1*L* sub-population of weak infectivity and S for strong infectivity. For the general analysis we simply consider periodic functions *K*(*x*), *μ*(*x*).

We distinguish between two relevant spatial scales, the scale of the individual patches *y*, represented by the inner squares and the global scale of the whole experiment *x*, represented by the checker board printing. Also, we assume that there is a small parameter *ε >* 0 such

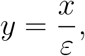

where *ε* represents the ratio between the local and global scales. In the experiment,

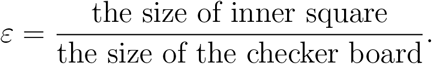

Thus, we can compute the *ε* for the 5 mm, 3 mm, and 1.5 mm inner squares as *ε* = 1/6, 0.1, 0.05, respectively.

We use standard assumptions in homogenization (see e.g. (Pavliotis and Stuart, 2008)) and assume that the growth rate *μ*(*y*) and the carrying capacity *K*(*y*) change only on the small scale *y* and they do not vary on the large scale *x*. The virus load depends on both scales, *v*(*x, y, t*) and the partial derivatives change as

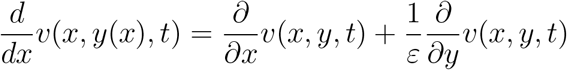

If we introduce this assumption into (1) we get the multiscale reaction-diffusion equation

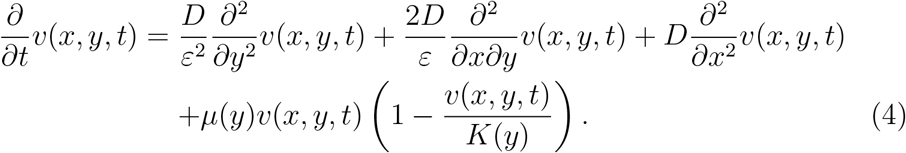

We are seeking here a leading order approximation (*v*_0_) to compute the viral load which is valid for small *ε.* To analyze this equation we use a perturbation expansion in the small parameter *ε* ≪ 1:

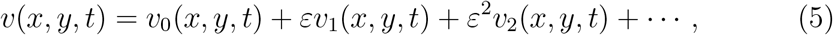

where all functions *v*_*j*_(*x, y, t*), *j* = 0, 1, 2, ... are assumed to be periodic in *y* of period 1.

We substitute this expansion (5) into (4) and collect terms of equal order in *ε*. The leading order term is of order *ε*^−2^:

- *ε*^−2^ : We obtain 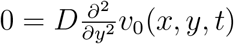, which leads to a general form

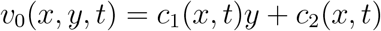 Since *v*_0_(*x, y, t*) is periodic in *y*, the first term *c*_1_ = 0 and we find that *v*_0_ does not depend on *y*. We write *v*_0_(*x, t*) instead of using *c*_2_(*x, t*).
- *ε*^−1^ : In this case we find

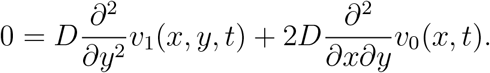 Since *v*_0_ does not depend on *y*, the second term is zero. Hence the first term is zero as well. Again arguing with periodicity, we find that also *v*1 is independent of *y* and we write *v*_1_(*x, t*).
- *ε*^0^ : Here we find

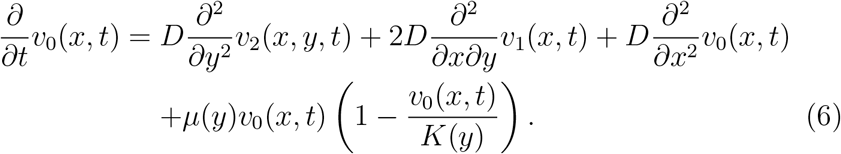 Instead of solving this equation for *v*_2_ we simply integrate over one period *y* ∈ [0, 1]: Since *v*_0_ and *v*_1_ do not depend on *y*, and since *v*_2_ is periodic, several terms simplify. We find the homogenized equation:

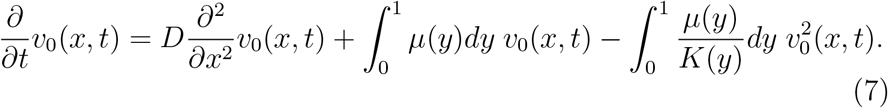

To understand (7) we introduce the *arithmetic mean* and the *harmonic mean* as

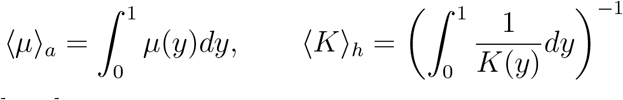

and we consider three cases:

#### Case 1

Consider *K*(*y*) = *K* constant. Then (7) becomes a standard Fisher-KPP equation

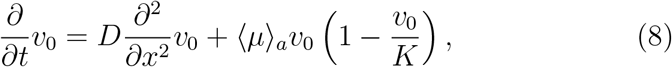

where the homogenized growth rate 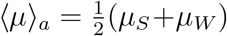 is the arithmetic mean of *μ*(*y*).

#### Case 2

Consider *μ*(*y*) = *μ* constant. In this case (7) becomes

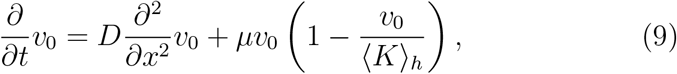

where the carrying capacity arises as harmonic mean of *K*(*y*):

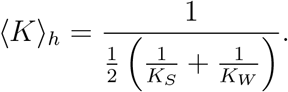

#### Case 3

We can also write the general homogenized equation (7) as a Fisher-KPP equation, however, with less intuitive average terms as

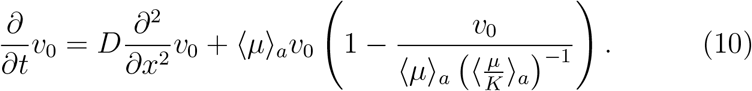

Here the effective growth rate and effective carrying capacity are

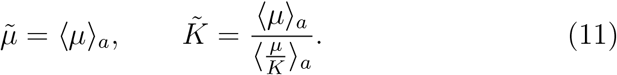

As illustrated in Figure 3, the fine printing of the virus hosts in patches of different sizes leads to different averaging. If the individual patches are large, then they can be considered as almost independent, and the growth rate and the carrying capacity will be the arithmetic means 〈*μ*〉_*a*_, 〈*K*〉_*a*_ of the patches. On the other hand, for finer and finer patches we have shown that the average growth rate is still the arithmetic mean 〈*μ*〉_*a*_, however, the carrying capacity uses the harmonic mean. For example in Case 2 above it is 〈*K*〉_*h*_ and it is known that

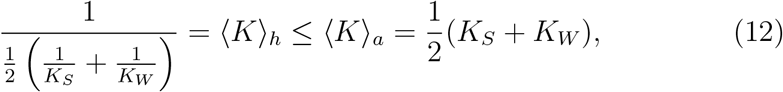

where equality is satisfied when *K*_*S*_ = *K*_*W*_. Hence a reduction of the overall carrying capacity for finer patches is a direct consequence of the averaging procedure.

**Figure 3:**
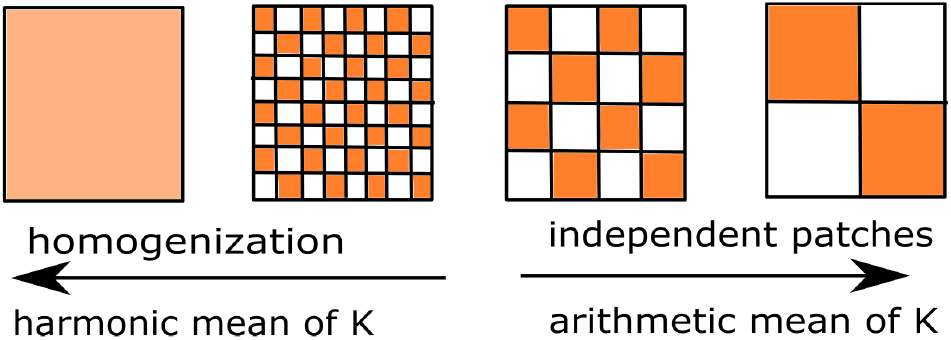
Sketch of a period patchy environment and the homogenization limit.

## 4. Application to the Fibroblast Experiments

Case 1, where *K*(*y*) =const., cannot describe the observed data, since the averaging of the growth rate does not change from coarse to fine experiments.

However, Cases 2 and 3 can. Since Case 2 is nested in Case 3, and since Case 2 is sufficient to explain the observed phenomenon, we focus our analysis on Case 2, where the virus growth rate *μ* is (almost) constant between the two cell types, while the carrying capacities are significantly different:

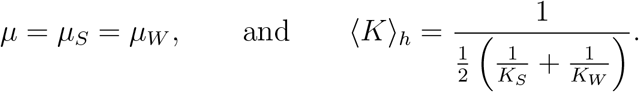

### 4.1. Fitting with PDEs

The above formulas (12) only give information about the mixed case and the most separated case. However, they do not give information about the intermediate scales such as the 1.5 mm and 3 mm experiments. To fit those data as well, we employ the Fisher-KPP model (1). Details of the numerical solution are given in Section 5.

We use a least-squares approach to estimate the error

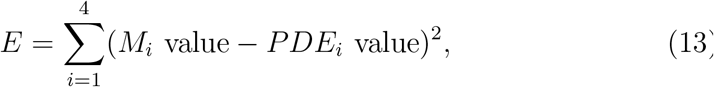

where i= 1,2,3,4 represent the inner squares sizes: mix, 1.5 mm, 3 mm, and 5 mm respectively. *M*_*i*_ denotes the measurement value and *PDE*_*i*_ the integral of the solution curve of the PDE.

### 4.2. Calibration I: Naive Approach

Based on the above calculations it is straight forward to simply compare the arithmetic means and harmonic means with the available data. In Table 1 we show the virus load data at *t* = 24*h* that correspond to the data shown in Figure 1 (B). The raw data are in Table A.7.

**Table 1:** Virus load data, standard deviation, standard error and error bars.

| Inner Square | Mix | 1.5 mm | 3 mm | 5mm |
| --- | --- | --- | --- | --- |
| Mean | 1.2957 | 1.3305 | 1.9304 | 3.3996 |
| Stand. Dev. | 0.2002 | 0.2917 | 0.8111 | 0.9968 |
| Stand. Error | 0.1156 | 0.1684 | 0.4683 | 0.5755 |
| Error Bar | [1.1801, 1.4113] | [1.1621, 1.4989] | [1.4621, 2.3986] | [2.8241, 3.9751] |

We assume for now that the 5 mm plate corresponds to the separated case, i.e. 〈*K*〉_*a*_ = 3.3996, while the mixed case corresponds to the harmonic mean 〈*K*〉_*h*_ = 1.2957. To find *K*_*S*_, *K*_*W*_ we then simply solve the two equations for the means (12) to find

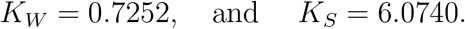

To investigate the agreement with the intermediate cases of 1.5 mm and 3 mm, we solve the full PDE (1). For this we also need the diffusion constant *D* and the virus growth rate *μ*. From the data in Tables A.5 and A.6, we found a range of values of the growth rate for the virus in weak and strong infectivity cells. While *μ*_*W*_ ∈ [0.1155, 0.3547], we found *μ*_*S*_ ∈ [0.1155, 0.4722]. Therefore, we choose the intermediate value 〈*μ*〉_*a*_ = 0.23 per hour.

We do not have any direct information from the data to estimate the value of the diffusion coefficient *D.* de Rioja et al. (2016) use a similar spatial virus model for cancer viral therapy, and they use a diffusion coefficient of *D*_Rioja_ = 0.0144 mm^2^ per hour. This corresponds to an invasion speed of *c* = 0.115 mm per hour, resulting in a distance travelled in 24h of 2.76 mm. This seems too small for our situation. If the travel distance is 2.76 mm in 24h, then the 3 mm and 5 mm cases would not have been able to effectively communicate viral load values over a time range of 24 h. Hence to explain the observed homogenization effect, we expect a significantly larger diffusion coefficient than 0.0144. Consequently, we take the diffusion coefficient as an unknown variable and compute the error (13) to the measurements. In Table 2 we vary *D* from 0.02 to 0.6 mm^2^ per hour and we solve the Fisher-KPP equation (1) as described in Section 5. We observe that for increasing *D* the fit for 1.5 mm gets better, the 3 mm fits well throughout, and the 5 mm fit gets worse. Hence in the end we do not observe a usable fit from this (naive) procedure. We show the data for the case of *D* = 0.5 as red curve in Figure 4 (A).

**Table 2:** Simulated virus load data when D is varied at t=24 hour.

| **D** | **0.02** | **0.05** | **0.1** | **0.3** | **0.5** | **0.6** | **Measurements** |
| --- | --- | --- | --- | --- | --- | --- | --- |
| Mix | 1.2365 | 1.2365 | 1.2365 | 1.2365 | 1.2365 | 1.2365 | $1.2957 \pm 0.1156$ |
| 1.5mm | 2.3060 | 2.0169 | 1.7582 | 1.4590 | 1.3800 | 1.3590 | $1.3305 \pm 0.1684$ |
| 3mm | 2.5734 | 2.4397 | 2.2762 | 1.9069 | 1.7304 | 1.6737 | $1.9304 \pm 0.4683$ |
| 5mm | 2.6757 | 2.6031 | 2.5175 | 2.2945 | 2.1431 | 2.0830 | $3.3996 \pm 0.5755$ |
| Fitting Data | No | No | No | No | No | No |  |

**Figure 4:**
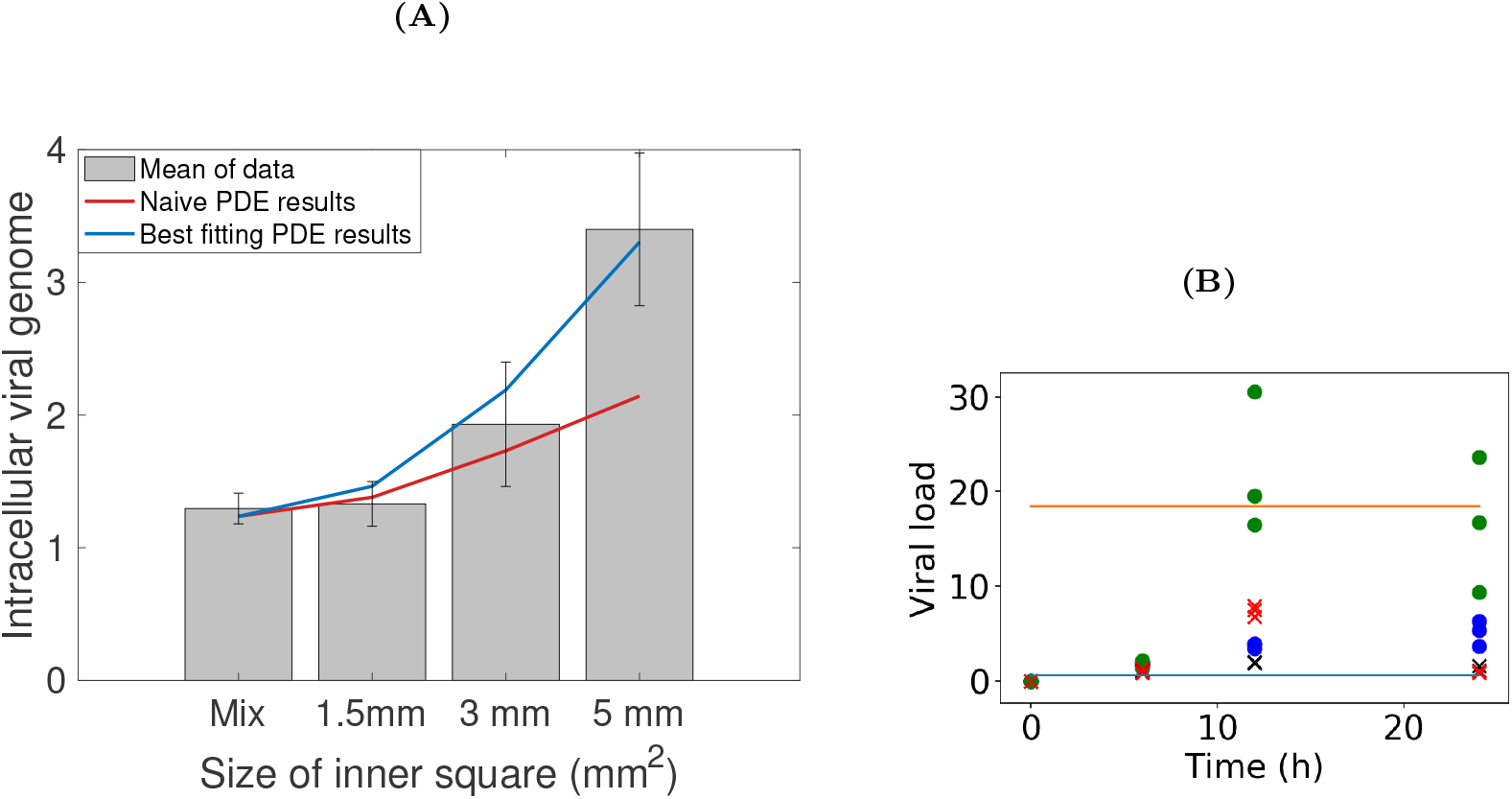
**(A):** Measurement values of virus load for the various checker board patterns. Overlaid is the naive fit for *D* = 0.5 in red and the best fit using a hypothetical 15 mm experiment in blue, **(B):** Comparison of the estimated carrying capacities *K*_*W*_, *K*_*S*_ with the virus load data of each cell type in isolation. The horizontal lines indicate the levels *K*_*W*_ = 0.6713 (blue), and *K*_*S*_ = 18.529 (orange). The data from Tables A.5 and A.6 are presented as (X,black)= weak cells experiment 1, (X,red)= weak cells experiment 2, (circle,blue)= strong cells experiment 1 and (circle,green)=strong cells experiment 2.

### 4.3. Calibration II: Extension to a 15 mm Case

The above mismatch for the 5 mm plate is related to the fact that the 5 mm squares are still relatively mixed, and they might not correspond to the fully separated state. Hence, numerically, we test this hypothesis by including a hypothetical 15 mm case, where the cell types are separated into one compartment for cells of weak infectivity and one compartment for cells of strong infectivity.

The corresponding virus load has not been measured (due to technical limitations of the bio-printing method), but we can still solve the PDE for this case. We define a maximal error tolerance of *E*_max_ = 0.1, which is the smallest error bar from the data (see Table 1). As we vary the values for *D* ∈ [0.02, 0.6] mm^2^ per hour and *K*_15*mm*_ ∈ [5.5, 12] we see in Figure (5 (A)) that the error is decreasing for increasing *D* and *K*_15*mm*_. The first value that is below the error tolerance of 0.1 is the choice of *D* = 0.5 (red marker in Figure 5 (A)). For larger values of *D* we still can decrease the error. A full minimization is, however, not very meaningful, since the fitting errors become much smaller than the measurement errors of the data. For fixed *D* = 0.5 we have a range of suitable values for *K*15*mm*. Here we can find a clear minimizer at *K*_15*mm*_ = 9.6 with an error of *E* = 0.0974. (see Figure 5 (B)). Hence for the purpose of our modelling we chose

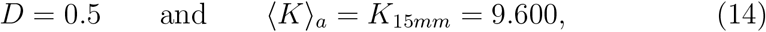

which, using (12) leads to

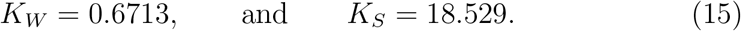

**Figure 5:**
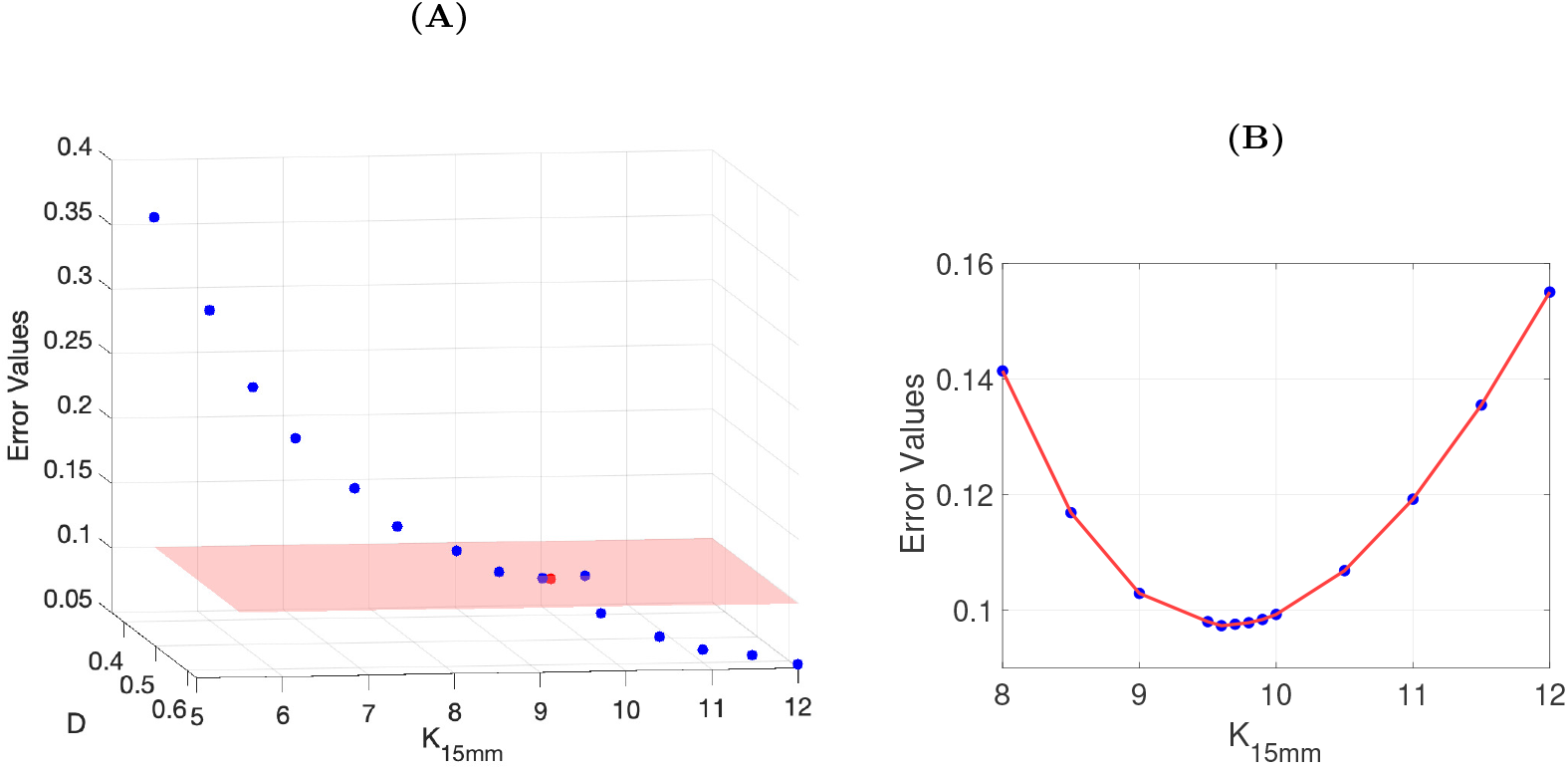
**(A)**: Error values at 24*h* when *K*_15*mm*_ and *D* are varied. The red marker indicates *D* = 0.5 and *K*_15*mm*_ = 9.6 which we chose as most suitable model parameter, **(B):** Optimization of *K*_15*mm*_ for fixed *D* = 0.5.

These results suggest that strong infectivity cells can support about 25 times more virus than weak infectivity cells.

In Table 3 and Figure 4 (A) we compare the measured values to the optimized PDE results and also record the error and relative error when *K*_15*mm*_ = 9.6 at t=24. We see that the model results are very close to the measurement, well within the error bars. A fit of this level of accuracy is quite uncommon for biological data, and we are confident that the chosen PDE model does explain the data well. We summarize the calibrated model parameters in Table 4.

**Table 3:**
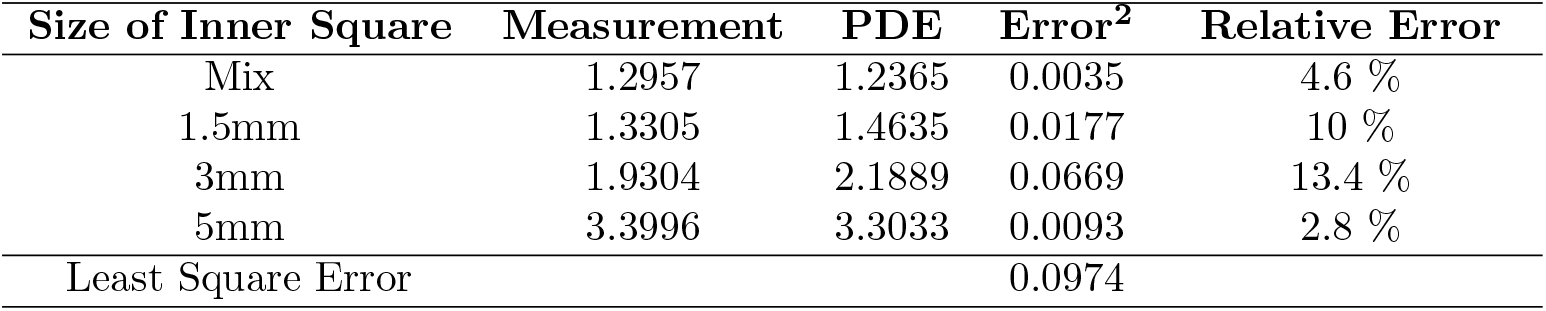
Comparison of the measurements with the optimized PDE model.

| Size of Inner Square | Measurement | PDE | Error<sup>2</sup> | Relative Error |
| --- | --- | --- | --- | --- |
| Mix | 1.2957 | 1.2365 | 0.0035 | 4.6 % |
| 1.5mm | 1.3305 | 1.4635 | 0.0177 | 10 % |
| 3mm | 1.9304 | 2.1889 | 0.0669 | 13.4 % |
| 5mm | 3.3996 | 3.3033 | 0.0093 | 2.8 % |
| Least Square Error |  |  | 0.0974 |  |

**Table 4:** Summary of calibrated model parameters.

| Parameter | Description | Value | Unit |
| --- | --- | --- | --- |
| $D$ | Diffusion coefficient | 0.5 | mm<sup>2</sup>per hour |
| $\langle\mu\rangle_a$ | Arith. mean growth of rate | 0.23 | per hour |
| $K_W$ | Carrying capacity of weak infectivity cells | 0.6713 | viral load |
| $K_S$ | Carrying capacity of strong infectivity cells | 18.529 | viral load |
| $K_{Mix} = \langle K \rangle_h$ | Carrying capacity of mixed plate | 1.2957 | viral load |
| $K_{15mm} = \langle K \rangle_a$ | Carrying capacity of 15 mm printed plate | 9.600 | viral load |

We further use our optimized PDE model to investigate the virus load after 12 hours as well. For the mix, 1.5 mm, 3 mm, and 5 mm we find simulated virus load numbers of 0.7372, 0.7643, 0.8183, and 0.8766, respectively. Although there is a slight increase from mixed to separated, the difference is small and would not be observable within measurement tolerances. Hence the homogenization effect would not be observed after 12h, an observation that has been confirmed in our experiments (experimental values not shown). The typical replication process of virus inside cells takes between 5 and 12 hours (Boianelli et al., 2015). Hence at 12h only a few cells would have released their virus contents, and the homogenization effect will not yet have kicked in.

### 4.4. Comparison to Experiments of Homogeneous Populations

We compare these results (15) with the experiments of viral load infections on each cell type in isolation. In Figure 4 (B) we plot the virus load data from tables A.5 and A.6 as functions of time. We use symbols and colors to distinguish between the cell types and experiments as (X, black)= weak cells experiment 1, (X, red)= weak cells experiment 2, (circle, blue)= strong cells experiment 1 and (circle, green)=strong cells experiment 2. In addition we plot the estimated *K*_*W*_ and *K*_*S*_ from (15) as horizontal lines.

We observe that there is a large difference in the virus load data between experiment 1 and 2 for each of the weak and strong cases. Hence the data do not seem to be directly comparable. The virus load of the strong responders is certainly one or two orders of magnitude larger than those of the weak responders, and our estimates reflect this fact nicely. However, the data do not allow a quantitative comparison.

## 5. Numerical Analysis of the PDE Model

The above results are based on numerical solutions of our PDE model (1), which we performed as follows.

### 5.1. Mix Plate

Since *μ*_*S*_ = *μ*_*W*_, we can solve the homogenized equation (9) for the Case 2 by a Forward-Time-Central-Space method (Smith, 1985). As *μ* and *K* are constant and the initial condition is non-negative, the solution should converge to the carrying capacity *K*_*Mix*_ = 〈*K*〉_*h*_ = 1.2957, which has been confirmed numerically as shown in Figure 6 (F).

### 5.2. Spatially Printed Plates

For the spatially printed plates, we consider two patch types, “strong” which represent strong infectivity cells and “weak” which represent weak infectivity cells. Accordingly, the carrying capacity is spatially constant within a patch but different between patches. While, the diffusion coefficient *D* and growth rate *μ* are the same in the two patches. We partition the entire interval into sub-intervals (’patches’) (*y*_*i*−1_, *y*_*i*_), *i* ∈ ℕ. Thus, we have

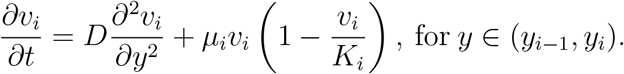

**Figure 6:**
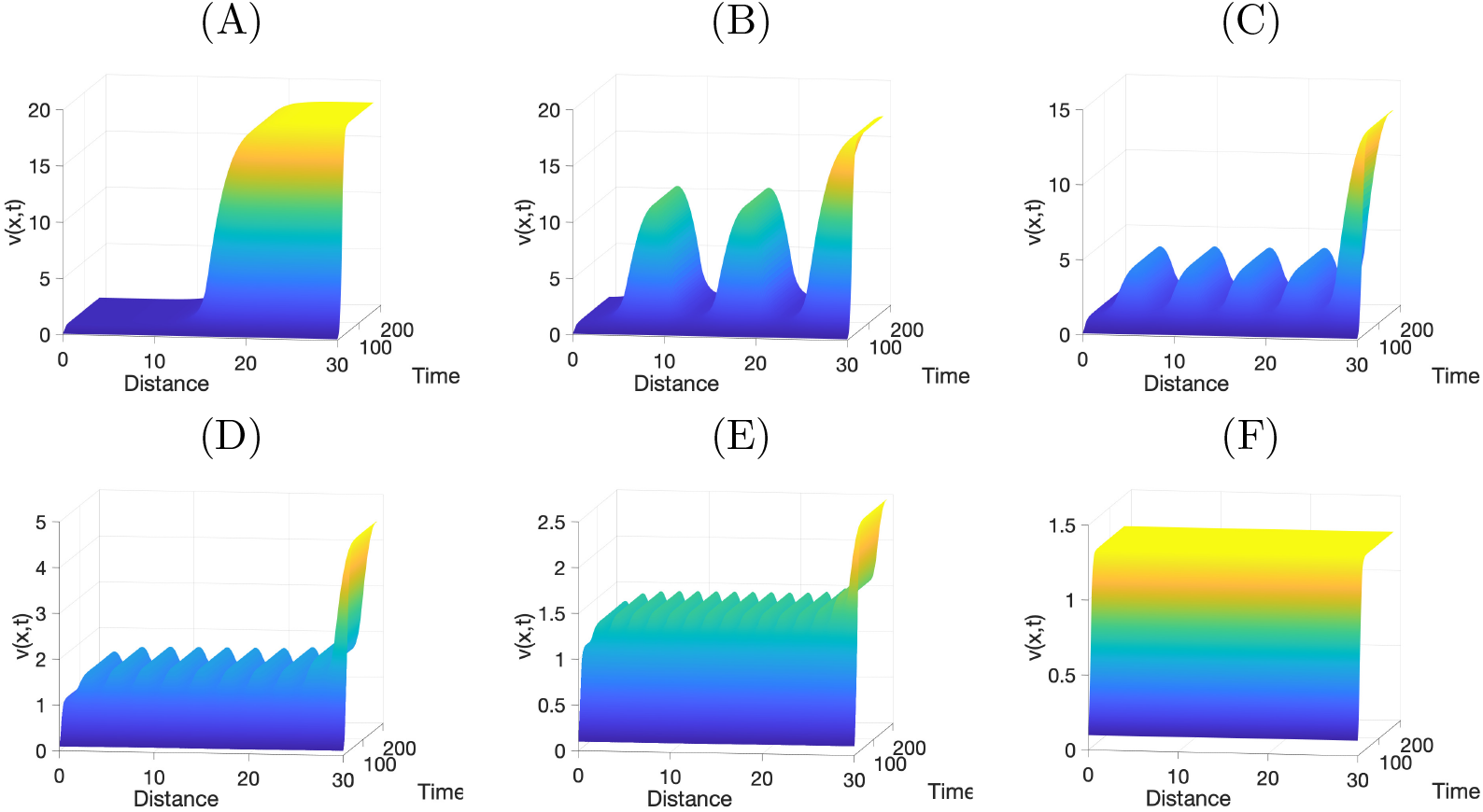
Numerical simulation when *μ*_*S*_ = *μ*_*W*_ = 0.23 and *D* = 0.5 with *v*(*x,* 0) = 0.1, and *v*_*x*_(0, *t*) = *v*_*x*_(30, *t*) = 0 for **(A):** 15 mm intervals, **(B):** 5 mm intervals, **(C):** 3 mm intervals, **(D):** 1.5 mm intervals, **(E):** 1 mm intervals, **(F):** mix plate. Note that the *z*-axis changes between these figures.

Since the diffusion coefficient does not vary between the patches, the flux is continuous across an interface

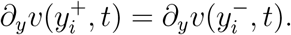

Here, 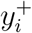 and 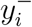 denote right and left sided limits at *y*_*i*_. The probability of a virion at interface *y*_*i*_ moving to the right or left is the same and equal to 0.5. Thus

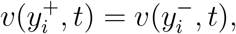

which ensures continuity of the solution at the interfaces.

The simulations in Figure 6 confirm our model assumption as only in the 15mm case (Figure 6 (A)) the carrying capacity of *K*_*S*_ = 18.529 of the strong infectivity cells is reached. In the other simulations (B)-(F), the inner maxima, corresponding to the strong cell type, are much lower than the carrying capacity of 18. We see the homogenization effect in action, as the relative differences between local maxima and local minima are flattened for decreasing inner interval sizes. We also observe an overshoot at the right boundary, which is due to the chosen Neumann boundary conditions.

## 6. Conclusion

In this paper, we apply the method of homogenization to a spatially structured Fisher-KPP model for the virus-load of a virus infection of cell cultures. The method of homogenization is well known in physics (Pavliotis and Stuart (2008)) and ecology (Yurk and Cobbold (2018); Garlick et al. (2011)), but here we apply it in a new way to microbiological data. We found a biological situation where the averaging matters. Where both, the arithmetic mean and the harmonic mean, provide valid information about viral load and they are not the same. It is the first experiment we know where the type of averaging matters for the infectivity of cell populations.

These observations have been made possible through the revolutionary technique of inkjet bioprinting as pioneered by Park et al (Park et al., 2017). Besides checker-board patterns, also other patterns, such as the Eiffel Tower for example, can be printed and analysed. This opens the door for modelling of more realistic heterogeneous tissue such as lung tissue for example.

The calibrated PDE model has been used to compute scenarios, which were not done experimentally, and hence serve as predictions, such as the 1 mm and 15 mm cases, as well as the homogenization effect after 12h. The viral load in the 1 mm inner square is reduced dramatically compared to the 5 mm and 15 mm and being very close to the measurement value of the mixed case. Also, using the formula for the invasion speed (2) we are able to compute the invasion speed in our experiment. For *D* = 0.5 and *μ* = 0.23 we find an invasion speed of 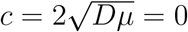 per hour. Which means in 24h virions travel a distance of 16.3 mm, which is about half of the domain size. Hence the travel distance within 24h is on the macroscopic scale of the experiment. This agrees well with the underlying assumption of homogenization that two spatial scales are considered.

Our observations are based on highly controlled experiments with two cell types and a clear geometric setup. Nevertheless, the effect of population mixing versus segregation on total viral load is likely to be present also in more natural occuring tissues. One such area would be viral therapy of cancer (Wodarz et al., 2012; de Rioja et al., 2016), where a heterogeneous cancer might react quite differently to viral therapy as compared to a homogeneous cancer. We found a 8 times increase of viral load from mixed to fully segregated. This can have a drastic impact on tissue response and resulting patient health. For example in a recent study by Néant et al. (2021) (Supplement Figure S1), on SARS-CoV-2 infection in France, a threshold of 10^6^ was identified as a predictor of COVID-19 mortality. A factor 8, as we found here, can easily make a difference in the infection outcome. The spatial distribution of SARS-CoV-2 virus in tissue has been studied in Getz et al. (2020), again, identifying to a highly heterogeneous lung tissue.

It should be noted that our model is spatially one dimensional, while the experiments are two dimensional. We argue that due to model symmetry a one-dimensional approach is sufficient. In fact, the model performs well, all results are within error tolerances, and we do not expect any further gain through a two dimensional version.

Still, there is always room for improvement. While we built the model exclusively for the available data, it would be worthwhile to include more of the viral infection dynamics, such as endocytosis, viral reproduction, exocytosis, as well as cell death (see Getz et al. (2020)). Also the transport of virions from cell to cell can be formulated in a much more detailed way, using cell membranes, fluid flow around the cells and possible “intracellular connections”, which are know to transport virions as well (Roberts et al., 2015). Here we get away with a simple effective diffusion process to explain the observed data.

Another way to evaluate the diffusion coefficient *D* is by applying the Stokes-Einstein equation for the diffusion coefficient *D* of a spherical particle of radius *r* in a fluid of dynamic viscosity *η* at absolute temperature *T* (Murray and Jackson, 1992), which in our case becomes

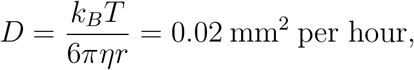

where *k*_*B*_ is Boltzmann’s constant, *r* = 50 *nm* is a typical virus size, and *η* = 0.00094 *pa. s* is the viscosity of DMEM (10 % FBS) medium at *T* = 25°*C* (Fröhlich et al., 2013). However, the simulation results show that this value is very low and it does not fit the data. We believe that a reason for this discrepancy is the formation of capillaries between cells that are touching along cell membranes. Breugem (2007) showed that depending on the capillary pore size, the diffusivity could increase by orders of magnitude. Again, in our case we have no information about the pore sizes between the cells in our experiments.

Biologically, our results are surprising because it means in viral infection, spatial separation between the cell sub-populations is not beneficially to the cell population as a whole. Therefore, for overall fitness of cell population, it is better to be mixed. We believe that this is a strategy the nature chooses to increase fitness.

## Data Availability

Data are in Appendix A

## Acknowledgements

AAB acknowledges support through United Arab Emirates University Scholarship. TH acknowledges support from the Natural Science and Engineering Research Council of Canada (NSERC) and a visiting professorship at the Korean Advanced Institute of Science and Technology (KAIST). AAB and TH thank the members of the Mathematical Biology Journal Club for their valuable comments.

## Appendix A. Experimental Data

Here we present the raw-data as measured in our viral load experiments. Table A.5 and A.6 represent the kinetics of influenza A virus separately in the homogeneous cell populations in a first and second experiment with triplicated data sets. In both experiments, the CT (HA) gene expression, CT (mGAPDH) gene expression, Δ*CT*, ΔΔ*CT* and RQ have been measured for weak and strong infectivity cells at t=0, t=6, t=12 and t=24 hours respectively. The viral load is represented by the RQ value, where the RQ is the relative quantification as we mentioned in section 2.

**Table A.5:** The growth rate of influenza A virus in weak and strong population in first experiment with three data sets for one day. N.D means not determined

| Sample name | CT (HA)<br>(Target gene) | CT (mGAPDH)<br>(Reference gene) | ΔCT | $\Delta\Delta$CT | RQ |
| --- | --- | --- | --- | --- | --- |
| Weak 0h-1 | 32.906 | 13.687 | N.D | N.D | N.D |
| Weak 6h-1 | 22.480 | 16.430 | 6.0500 | 0.1090 | 0.9272 |
| Weak 12h-1 | 20.533 | 15.500 | 5.0330 | -0.9080 | 1.8764 |
| Weak 24h-1 | 20.008 | 14.719 | 5.2890 | -0.6520 | 1.5713 |
| Weak 0h-2 | 33.670 | 14.912 | N.D | N.D | N.D |
| Weak 6h-2 | 22.681 | 16.685 | 5.9960 | 0.0550 | 0.9626 |
| Weak 12h-2 | 20.690 | 15.757 | 4.9330 | -1.0080 | 2.0111 |
| Weak 24h-2 | 20.184 | 14.417 | 5.7670 | -0.1740 | 1.1282 |
| Weak 0h-3 | 33.175 | 15.184 | N.D | N.D | N.D |
| Weak 6h-3 | 22.137 | 16.360 | 5.7770 | -0.1640 | 1.1204 |
| Weak 12h-3 | 20.611 | 15.687 | 4.9240 | -1.0170 | 2.0237 |
| Weak 24h-3 | 19.864 | 13.947 | 5.9170 | -0.0240 | 1.0168 |
| Strong 0h-1 | 33.714 | 14.757 | N.D | N.D | N.D |
| Strong 6h-1 | 21.071 | 15.869 | 5.2020 | -0.7390 | 1.6690 |
| Strong 12h-1 | 9.745 | 15.780 | 3.9650 | -1.9760 | 3.9340 |
| Strong 24h-1 | 17.433 | 13.385 | 4.0480 | -1.8930 | 3.7141 |
| Strong 0h-2 | 34.002 | 13.825 | N.D | N.D | N.D |
| Strong 6h-2 | 20.420 | 15.398 | 5.0220 | -0.9190 | 1.8908 |
| Strong 12h-2 | 19.875 | 15.901 | 3.9740 | -1.9670 | 3.9095 |
| Strong 24h-2 | 18.039 | 14.757 | 3.2820 | -2.6590 | 6.3160 |
| Strong 0h-3 | 34.402 | 14.588 | N.D | N.D | N.D |
| Strong 6h-3 | 20.254 | 15.299 | 4.9550 | -0.9860 | 1.9807 |
| Strong 12h-3 | 19.902 | 15.728 | 4.1740 | -1.7670 | 3.4035 |
| Strong 24h-3 | 18.447 | 14.939 | 3.5080 | -2.4330 | 5.4002 |

**Table A.6:** The growth rate of influenza A virus in weak and strong population in second experiment with three data sets for one day. N.D means not determined

| Sample name | CT (HA)<br>(Target gene) | CT (mGAPDH)<br>(Reference gene) | ΔCT | $\Delta\Delta$CT | RQ |
| --- | --- | --- | --- | --- | --- |
| Weak 0h-1 | N.D | 13.927 | N.D | N.D | N.D |
| Weak 6h-1 | 24.539 | 19.245 | 5.2940 | -0.3600 | 1.2834 |
| Weak 12h-1 | 18.613 | 15.948 | 2.6650 | -2.9890 | 7.9390 |
| Weak 24h-1 | 20.678 | 14.778 | 5.9000 | 0.2460 | 0.8432 |
| Weak 0h-2 | N.D | 14.059 | N.D | N.D | N.D |
| Weak 6h-2 | 23.796 | 17.951 | 5.8450 | 0.1910 | 0.8760 |
| Weak 12h-2 | 19.146 | 16.261 | 2.8850 | -2.7690 | 6.8164 |
| Weak 24h-2 | 20.581 | 15.002 | 5.5790 | -0.0750 | 1.0534 |
| Weak 0h-3 | 37.975 | 14.316 | N.D | N.D | N.D |
| Weak 6h-3 | 23.748 | 17.925 | 5.8230 | 0.1690 | 0.8895 |
| Weak 12h-3 | 19.805 | 17.057 | 2.7480 | -2.9060 | 7.4950 |
| Weak 24h-3 | 20.570 | 14.960 | 5.6100 | -0.0440 | 1.0310 |
| Strong 0h-1 | N.D | 14.942 | N.D | N.D | N.D |
| Strong 6h-1 | 21.493 | 16.308 | 5.1850 | -0.4690 | 1.3841 |
| Strong 12h-1 | 16.076 | 14.713 | 1.3630 | -4.2910 | 19.576 |
| Strong 24h-1 | 15.642 | 14.052 | 1.5900 | -4.0640 | 16.726 |
| Strong 0h-2 | 36.907 | 14.285 | N.D | N.D | N.D |
| Strong 6h-2 | 21.513 | 16.950 | 4.5630 | -1.0910 | 2.1302 |
| Strong 12h-2 | 16.607 | 15.885 | 0.7220 | -4.9320 | 30.527 |
| Strong 24h-2 | 16.010 | 13.580 | 2.4300 | -3.2240 | 9.3437 |
| Strong 0h-3 | N.D | 13.836 | N.D | N.D | N.D |
| Strong 6h-3 | 20.211 | 15.401 | 4.8100 | -0.8440 | 1.7950 |
| Strong 12h-3 | 15.965 | 14.355 | 1.6100 | -4.0440 | 16.495 |
| Strong 24h-3 | 15.060 | 13.967 | 1.0930 | -4.5610 | 23.605 |

Table A.7, represents the viral load for the checker board experiments at one day with three independent samples. In the micro-pattering experiment, the CT (HA) gene expression, CT (GAPDH) gene expression, RQ and the average RQ (average viral load) have been measured in the mix plate, 1.5 mm, 3 mm and 5 mm, respectively.

**Table A.7:** The influenza A viral load on the micro-patterning for the different inner square sizes with three samples at one day.

| Size of inner square | CT (HA) | CT (GAPDH) | RQ | Average RQ |
| --- | --- | --- | --- | --- |
| Mix-1 | 15.366 | 15.157 | 1.0669 | 1.2957 |
| Mix-2 | 14.849 | 15.013 | 1.3816 |  |
| Mix-3 | 14.695 | 14.917 | 1.4387 |  |
| 1.5 mm-1 | 15.148 | 14.845 | 1.0000 | 1.3305 |
| 1.5 mm-2 | 15.172 | 15.395 | 1.4396 |  |
| 1.5 mm-3 | 15.461 | 15.793 | 1.5520 |  |
| 3 mm-1 | 14.621 | 15.638 | 1.7034 | 1.9304 |
| 3 mm-2 | 15.266 | 15.087 | 1.2569 |  |
| 3 mm-3 | 14.440 | 15.294 | 2.8308 |  |
| 5 mm-1 | 14.191 | 15.060 | 2.2519 | 3.3996 |
| 5 mm-2 | 14.948 | 16.609 | 3.8978 |  |
| 5 mm-3 | 14.547 | 16.262 | 4.0492 |  |

## Notes

### Competing Interest Statement

The authors have declared no competing interest.

